## Supplemental Table for "Oxidized low-density lipoprotein potentiates angiotensin II-induced Gq activation through the AT1-LOX1 receptor complex: Implications for renal dysfunction"

Supplemental Table. Primer Sequences used in this study

| gene (rat) | Forward (5`-3`) | Reverse (5`-3`) |
|---|---|---|
| *GAPDH* | AGGAGTAAGAAACCCTGGAC | CTGGGATGGAATTGTGAG |
| *AT1a* | ACAACTGCCTGAACCCTCTG | CCTCAAAACAAGACGCAGGC |
| *LOX1* | AGAAGCCTAAAGGGCTGCAT | ACATCTGCCCCTCCAGGATA |
| *TGFβ1* | AGGAGGAATTTGGCCAGGTG | GCTCACGAGGAGGCTAATCC |
| *TNFα* | GGCATGGATCTCAAAGACAACC | AAATCGGCTGACGGTGTGG |
| *IL-1β* | CACCTCTCAAGCAGAGCACAG | GGGTTCCATGGTGAAGTCAAC |
| *IL-6* | GTCAACTCCATCTGCCCTTCAG | GGCAGTGGCTGTCAACAACAT |
| *MCP-1* | AGGCAGATGCAGTTAATGCCC | ACACCTGCTGCTGGTGATTCTC |
| *ICAM-1* | AGCTCTTCAAGCTGAGCGACAT | ACTCGCTCTGGGAACGAATACA |
| *VCAM-1* | GCGAAGGAAACTGGAGAAGACA | ACACATTAGGGACCGTGCAGTT |
| *P22phox* | TCCACTTACTGCTGTCCGT | TCAATGGGAGTCCACTGCT |
| *P67phox* | AGCAGAAGAGCAGTTAGCATTGG | TGCTTTCCATGGCCTTGTC |
| *P91phox* | TGGTGATGTTAGTGGGAGC | CTTTCTTGCATCTGGGTCT |
| *fibronectin* | AAAGCCAGCCCCTGGTTC | AGGTCACCTGTACCTGGAA |
| *Col1a* | GACATCCCTGAAGTCAGCTGC | TCCCTTGGGTCCCTCGAC |
| *Col4a* | TGGCCTTGGAGGAAACTTTG | CTTGGAAACCTTGTGGACCAG |

| gene (mouse) | Forward (5`-3`) | Reverse (5`-3`) |
|---|---|---|
| *GAPDH* | TTCCATCCTCCAGAAACCAG | CTCAGACCCCAGATCCAGAA |
| *AT1a* | TGTCTGGCCGGAGAGGACT | TCTTTCATATGTTAAGTCCGGGAGA |
| *LOX1* | GGCCAACCATGGCTATGGGAGAATGG | CAGCGAACACAGCTCCGTCTTGAAGG |
| *TGFβ1* | TGATACGCCTGAGTGGCTGTCT | CACAAGAGCAGTGAGCGCTGAA |
| *TNFα* | GGCTGCCCCGACTACGT | ACTTTCTCCTGGTATGAGATAGCAAAT |
| *IL-1β* | GTCACAAGAAACCATGGCACAT | GCCCATCAGAGGCAAGGA |
| *IL-6* | CTGCAAGAGACTTCCATCCAGTT | AGGGAAGGCCGTGGTTGT |
| *MCP-1* | AAAACAGCATACATGGGAGACT | ATCCAGGGCACATATGCAGAG |
| *ICAM-1* | GTGATGCTCAGGTATCCATCCA | CACAGTTCTCAAAGCACAGCG |
| *VCAM-1* | AGTTGGGGATTCGGTTGTTCT | CCCCTCATTCCTTACCACCC |
| *P22phox* | GTCCACCATGGAGCGATGTG | CAATGGCCAAGCAGACGGTC |
| *P67phox* | CTGGCTGAGGCCATCAGACT | AGGCCACTGCAGAGTGCTTG |
| *P91phox* | TTGGGTCAGCACTGGCTCTG | TGGCGGTGTGCAGTGCTATC |
| *fibronectin* | AAGACCATACCTGCCGAATG | GAACATGACCGATTTGGACC |
| *Col1a* | GAAGCACGTCTGGTTTGGA | ACTCGAACGGGAATCCATC |
| *Col1b* | CCAACAAGCATGTCTGGTTAGGA | TCAAACTGGCTGCCACCAT |
| *Col3a* | GAAAGAGGATCTGAGGGCTCG | GGGTGAAAAGCCACCAGACT |
| *Col4a* | CCAAAAGGACAGCAAGGTGTG | CCAGGAAACCCTCTTGGACC |
| *P40phox* | GCCGCTATCGCCAGTTCTAC | GCAGGCTCAGGAGGTTCTTC |
| *P47phox* | GATGTTCCCCATTGAGGCCG | GTTTCAGGTCATCAGGCCGC |
| *AT1b* | GAGACCAGACAAGACACGCA | GTGAATTCAAAATGCACCCGT |
| *α-SMA* | GACGTACAACTGGTATTGTG | TCAGGATCTTCATGAGGTAG |
| *Vimentin* | AGCTGCTAACTACCAGGACACTATTG | CGAAGGTGACGAGCCATCTC |
| *E-cadherin* | GTCTCCTCATGGCTTTGC | CTTTAGATGCCGCTTCAC |
| *Cadherin-16* | CTGGCAGCGATAGGCTTCAT | TGGGGCTGCTTGGATCATTC |
| *Kim-1* | GCTGCAATGGAGATGCCAAC | TCCTCAGATGACCCTTGGGA |
| *NAGL* | TCTGTCCCCACCGACCAA | GGAAAGATGGAGTGGCAGACA |
| *COX2* | TGGTGAAAACTGTACTACACCTGA | CTTCGCAGGAAGGGGATGTT |
| *VEGF-a* | CTTGTTCAGAGCGGAGAAAGC | ACATCTGCAAGTACGTTCGTT |
| *Podocin* | GTGTCCAAAGCCATCCAGTT | GTCTTTGTGCCTCAGCTTCC |
| *α-actinin4* | GCCATCCAGGACATCTCTGT | CCGCAGCTTGTCATACTCAA |
| *CD2AP* | AGGAATTCAGCCACATCCAC | TTGAGGGAAACAGTCCCAAC |
